# Reliable estimates of Japanese encephalitis virus evolutionary rate supported by a formal test of temporal signal

**DOI:** 10.1101/2024.03.27.586903

**Authors:** Polina Ukraintceva, Artem Bondaryuk, Rodion Sakhabeev

## Abstract

The single-stranded RNA viruses exhibit stupendously high evolutionary rate spanned several orders of magnitude. The accurate and reliable estimates of the virus evolutionary rate are of importance for the robust interpretation of outbreak investigations and are critical for understanding short-term virus transmission patterns. In the previous studies, the substitution rate of the Japanese encephalitis virus (JEV) and time to the most recent common ancestor (MRCA) were reported, however, the estimates are substantially different between some of the studies. Importantly, the temporal signal in data was not evaluated by a formal test. The purpose of our study is to estimate the substitution rate and the age of the JEV common clade (GI-GV) and two JEV genotypes (GI and GIII) supported by the formal Bayesian analysis of temporal signal in the open reading frame (ORF) nucleotide data. Additionally, we assessed temporal signal in the data from the previous work to explain the observed discrepancy between the reported JEV evolutionary rate estimates. Results showed that the substitution rate of the JEV GI-GV was 2.41×10^−4^ nucleotide substitution per site per year (s/s/y) with 95% highest posterior density (HPD) interval of 1.67×10^−4^–3.14×10^−4^ or approximately 2.5 nucleotide substitutions per ORF (10,296 nucleotides) per year. That is one of the lowest substitution rates among the species closely related to JEV and other well-studied members of mosquito-borne flaviviruses. The substitution rate of GI and GIII was evaluated to be 4.13×10^−4^ s/s/y (95% HPD interval, 3.45×10^−4^– 4.82×10^−4^) and 6.17×10^−5^ s/s/y (95% HPD interval 3.84×10^−5^–8.62×10^−5^), accordingly, with the ages of clades of 153 (95% HPD, 87–237) and 216 (95% HPD, 139–317), respectively. The mean root height of JEV is 1234 years (95% HPD, 407–2333).

## 1. Introduction

Japanese encephalitis virus (JEV) belongs to the species *Orthoflavivirus japonicum,* family *Flaviviridae* (Postler et al. 2023). JEV is the member of the mosquito-borne flavivirus (MBFV) group. The JEV genome consists of single-stranded positive-sense RNA with a length of about 10,965 nucleotides (nt). The genome encompasses a 10,296 nt single open reading frame (ORF), encoding a polyprotein, which includes three structural proteins: capsid (C), precursor membrane (prM) and envelope (E) proteins, and seven non-structural ones: NS1, NS2a, NS2b, NS3, N4a, NS4b, NS5 (Sumiyoshi et al. 1987; Xu et al. 2023).

Mosquitoes are the vectors for JEV, in particular, *Culex tritaeniorhynchus* and *Culex annulirostris*, which were recognized as principal ones (Van den Eynde et al. 2022). The virus is zoonotic, and it is transmitted in most cases between pigs or birds by mosquitoes, however, Ricklin et al. (2016) described the direct way of virus transmission between two pigs via oronasal secretions. Humans, cattle and horses are dead-end hosts, due to insufficient viremia to infect mosquitoes. When these hosts are infected, encephalitis can develop with fever, convulsions, coma and even death (Van den Eynde et al. 2022), (Ladreyt et al. 2019).

Approximately 3 billion people live in JEV epidemic areas according to the World Health Organization data (https://www.who.int/news-room/fact-sheets/detail/japanese-encephalitis, Retrieved 28 March 2024). JEV is the major causative agent of human virus encephalitis disease in Asia. Now the virus is endemic in the Torres Strait region of Australia and Asia, including 24 countries, namely China, Korea, Indonesia, Malaysia, Japan and even Pakistan (Van den Eynde et al. 2022), (Mackenzie 2005). Additionally, the virus was observed outside the endemic areas in Italy (however, the data of the study are limited with one short sequence, wherein JEV has not been registered in Europe since then) (Ravanini et al. 2012) and Angola (Simon-Loriere et al. 2017).

Today, between 30,000 (Van den Eynde et al. 2022) and 69,000 (Auerswald et al., 2021) cases registered annually according to different sources, but it is considered as underestimated values due to insufficient monitoring and diagnosis. Despite the fact that effective and safe vaccines have been developed (Vannice et al. 2021), JEV is still one of the significant agents causing viral encephalitis in the territory of Asia. Thus, to monitor and control this disease, it is important to scrutinize evolutionary driving forces, which determine virus spread, genetic variability and pathogenicity.

While recombination has previously been shown to contribute to JEV evolution (Sistrom et al. 2024), most research focuses on the substitution rate. The single-stranded RNA viruses exhibit stupendously high evolutionary rate spanned several orders of magnitude (Duffy et al. 2008; Duchene et al. 2014). The members of MBFVs evolve at a rate in the order of 10^−4^ nucleotide substitutions per site per year (s/s/y) (Delatorre et al. 2019; Liu et al. 2019; Srihi et al. 2021; Zecchin et al. 2021; Stica et al. 2022; Xu et al. 2023). The evolutionary rate of JEV has been assessed several times but the estimates vary widely in the different sources spanning two orders of magnitude (10^−5^ to 10^−4^ s/s/y) (Mohammed et al. 2011; Pan et al. 2011; Gao et al. 2015; Li et al. 2023; Xu et al. 2023). It is important to note that none of the previous works has performed formal or any evaluation of temporal signal in data sets prior to molecular clock application, and, therefore, the reliability of the reported estimates is questionable. The accurate estimates of the evolutionary rate are of importance for the robust interpretation of outbreak investigations and are critical for understanding short-term virus transmission patterns (Holmes et al. 2016; Theze et al. 2018; O’Toole et al. 2023; Crits-Christoph et al. 2024). Besides, inferring evolutionary timescales helps to describe and quantify the population dynamic processes that gave rise to the virus genetic diversity observed.

The aim of our study is the reliable assessment of substitution rates and ages of JEV with its genotypes and the (re-)evaluation of temporal signal in the data sets from the previous studies using the Bayesian phylogenetic approach. To assess JEV substitution rate, we used 325 complete ORF nucleotide sequences of all five JEV genotypes with known collection dates available at the start of a study (January 2023). Evolutionary rate and divergence times obtained were compared with the previous estimates of JEV evolutionary rate and the rate of virus species, closely related to JEV, and other well-studied MBFVs. Bayesian evaluation of temporal signal makes reported results the first formally tested assessment of JEV substitution rate.

## 2. Materials and methods

### 2.1 Nucleotide data set

As of January 2023, 440 nucleotide sequences of complete or near complete genomes of JEV were found in GenBank using BLAST (http://blast.ncbi.nlm.nih.gov/Blast.cgi). Clones, vaccine and chimeric strains were removed from the data set and then collection dates were identified for 367 strains. Recombinant sequences detected by RDP5 (Martin et al. 2021) were removed from the alignment. Thus, our research was conducted using 325 nucleotide sequences of a polyprotein gene.

Next, sequence alignment was done with MAFFT v.7.490 (Katoh and Standley 2013). After that, the alignment was visualized in AliView v.1.24 (Larsson 2014) to exclude ORFs with stop-codons. Non-coding regions were cut off using the NCBI reference sequences (NC_001437.1).

JEV genotypes were determined in preliminary phylogenetic analysis in IQ-TREE v.1.6.12 using the tree topology reported by (Takhampunya et al. 2011). It was revealed that 134 samples belong to GI, 7 to GII, 171 to GIII, 6 to GIV and 7 to GV. Thus, only GI and GIII have a sufficient number of sequences to obtain reliable statistics in the further independent Bayesian phylogenetic analyses. Therefore, we subsequently analyzed three data sets—the common clade of JEV (JEV GI-GV), GI and GIII.

NCBI accession numbers, as well as fasta and BEAST xml files can be found in the supplementary materials.

### 2.2 Temporal signal assessment

Temporal signal (or structure) assessment is the key step in substitution rate evaluation. In the case of absence of temporal signal, the program (BEAST in our case) inferred values based on the priors (e.g., population size prior) but not the data that potentially bias the entire analysis.

Temporal structure in each of the data sets (JEV GI-GV, GI and GIII) was assessed with the Bayesian evaluation of temporal signal (BETS) (Duchene et al. 2020) in BEAST v.2.6.7.

BETS involves the comparison of time-stamped (heterochronous) and isochronous models (M_het_ and M_iso_) based on the difference between log marginal likelihoods (for a tutorial of using BETS see: https://beast.community/bets_tutorial). The log marginal likelihoods for each model were calculated using the path sampling estimator implemented in the MODEL_SELECTION package from the BEAST2 Package Manager. The number of steps was set as 200 and the lengths of MCMC for each step were determined so that the product of the number of steps and MCMC length equaled the length of the MCMC during the main phylogenetic analysis in BEAST with an effective sample size (ESS) at least of 200. Priors on model parameters (molecular clock, substitution model, population model) were chosen similar to those used in the main analysis (Table S1) with the exception of the tree prior—in this case, we replaced the Bayesian skyline with the coalescent constant population with a proper prior (i.e., integrated to 1) on effective population size (Table 1).

**Table 1.** Priors used in the Bayesian evaluation of temporal signal (BETS) in all three data sets (JEV GI-GV, GI, GIII)

| Model | Parameter |  | Prior |
| --- | --- | --- | --- |
| Strict clock | Clock rate | | $\Gamma(\alpha = 0.1, \beta = 100)$ |
| Optimized relaxed clock | Relative branch rates | | LogNormal($M = 1.0, S = 0.2$) |
| | Sigma | | $\Gamma(\alpha = 0.5396, \beta = 0.3819)$ |
| | Mean clock rate | | $\Gamma(\alpha = 0.1, \beta = 100)$ |
| Rate heterogeneity | Shape of $\Gamma$ distribution | | Exp($M = 1.0$) |
|  | Proportion of invariant sites |  | Uniform(0.0, 1.0) |
| Substitution model | GTR rates | Transitions | $\Gamma(\alpha = 0.05, \beta = 20)$ |
| | | Transversions | $\Gamma(\alpha = 0.05, \beta = 10)$ |
| Tree prior (coalescent constant size) | Effective population size | | Exp($M = 100$) |

The comparison of different molecular clock can also be involved in BETS for calculation of marginal likelihoods for each model. Therefore, we included a strict clock (SC) and an optimized relaxed clock (ORC) (Douglas et al. 2021) models into the comparison.

The difference between log marginal likelihoods of competitive models (i.e., log Bayes factor) of at least 5.0 indicates very strong support, a value of 3 means strong support and a value of 1 is considered to be positive evidence (Kass and Raftery 1995).

For comparison purposes, we conducted widely used root-to-tip (RTT) regression, which is an informal test with well-known limitations: data points are not independent and strict clocklike behavior is assumed. RTT statistics were computed with Clockor2 (Featherstone et al. 2024). The trees for Clockor2 were reconstructed in IQ-TREE v.1.6.12 with the best-fit model determined for each data set using ModelFinder (Kalyaanamoorthy et al. 2017). The JEV GI-GV tree was rooted in Clockor2 by maximizing the coefficient of determination (R^2^) values. To calculate R^2^ for JEV GI and GIII genotypes, the trees were rooted using information from the JEV GI-GV phylogeny.

Additionally, temporal signal was evaluated using BETS (path sampling method) in the data sets from the previous works of Gao et al. (2015), Li et al. (2023), Mohammed et al. (2011), Xu et al. (2023), and Pan et al. (2011) available as of November 2023. In total, 9 data sets were gathered (Table 2):

**Table 2.** Data sets of JEV open reading frame sequences from the previous studies.

| Study | JEV genotype | Number of samples | Sampling window (years) |
| --- | --- | --- | --- |
| Mohammed et al. (2011) | GI-GV | 34 | 74.4 |
|  | GI | 10 | 15.4 |
|  | GIII | 21 | 73.0 |
| Pan et al. (2011) | GI-GV | 95 | 74.0 |
|  | GI | 43 | 32.0 |
|  | GIII | 50 | 70.6 |
| Gao et al. (2015) | GI-GV | 98 | 74.0 |
| Li et al. (2023) | GI | 131 | 42.0 |
| Xu et al. (2023) | GI-GV | 115 | 87.1 |

One sample from the study of Mohammed et al. (2011) and three samples from the study of Pan et al. (2011) previously genotyped as GIII are absent in our data sets since these sequences were removed from GenBank at the request of the authors.

As a molecular clock model, we used an uncorrelated log-normal relaxed clock (UCLD) as in the original works.

### 2.3 Dated phylogeny reconstruction and substitution rate evaluation

Bayesian phylogenetic analysis using JEV GI-GV, GI and GIII data sets was performed in BEAST v.2.6.7 (Bouckaert et al. 2019). The selection of substitution model was based on the BIC previously assessed by ModelFinder in IQ-TREE (GTR+Γ_4_+I for all data sets). Molecular clock models were compared using the log marginal likelihoods calculated during the BETS step. As a tree prior, we specified a flexible non-parametric Bayesian Skyline model. For details on all priors, see the supplementary.

The lengths of MCMC in each run were chosen so that the ESS associated with estimated parameters reached at least 200 samples. The inspection of model parameters and MCMC convergence was carried out in Tracer v.1.7.1 (Rambaut et al. 2018). The burn-in thresholds of MCMC chains were defined individually depending on the chain behavior. Maximum clade credible (MCC) trees were annotated in TreeAnnotator (Bouckaert et al. 2019). Due to high computational cost, BEAST was run using a BEAGLE library (Ayres et al. 2012) on a GPU (NVIDIA GeForce RTX 3070).

### 2.4 Comparing evolutionary rates among JEV subtypes estimated using a relaxed molecular clock

Relaxed molecular clocks allow evolutionary rates vary among tree branches. Therefore, the among-branch rate variation should be taken into account when comparing the substitution rates of different JEV subtypes. To address this problem, we calculated the posterior probability for a hypothesis that a random rate value from a lognormal rate distribution of one genotype is larger than a random rate value from a lognormal rate distribution of another genotype. To approximate this probability, we performed Monte Carlo simulations in R (Figure 2):

**Figure 1.**
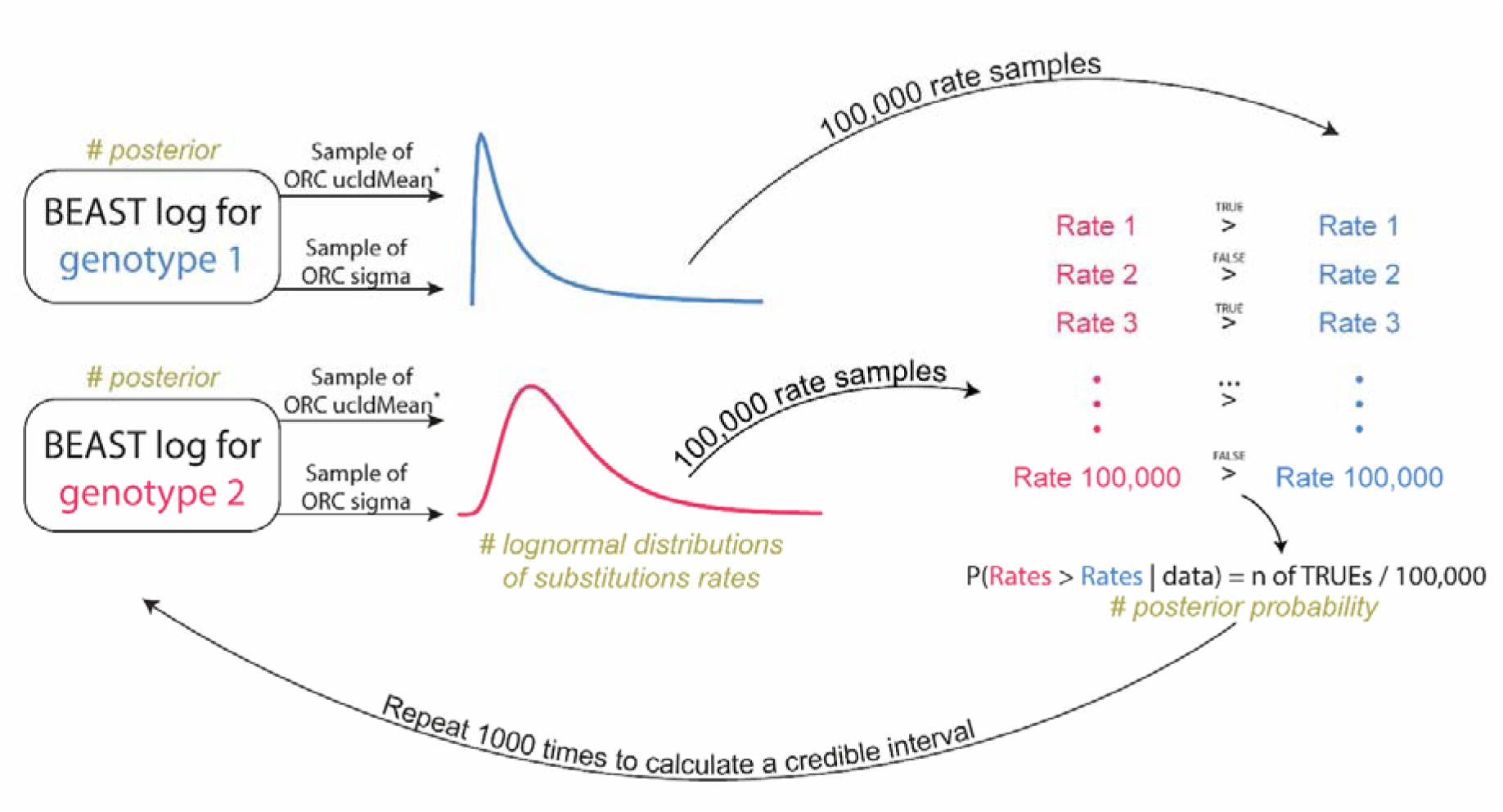
Flow chart of the comparative analysis of substitution rates between a pair of virus genotypes evaluated using relaxed molecular clock. Random samples from lognormal distributions were drawn with Monte Carlo simulations in R. Asterix—ucldMean parameter values in real space must be transformed to log space: 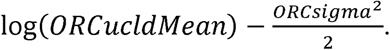

**Figure 2.**
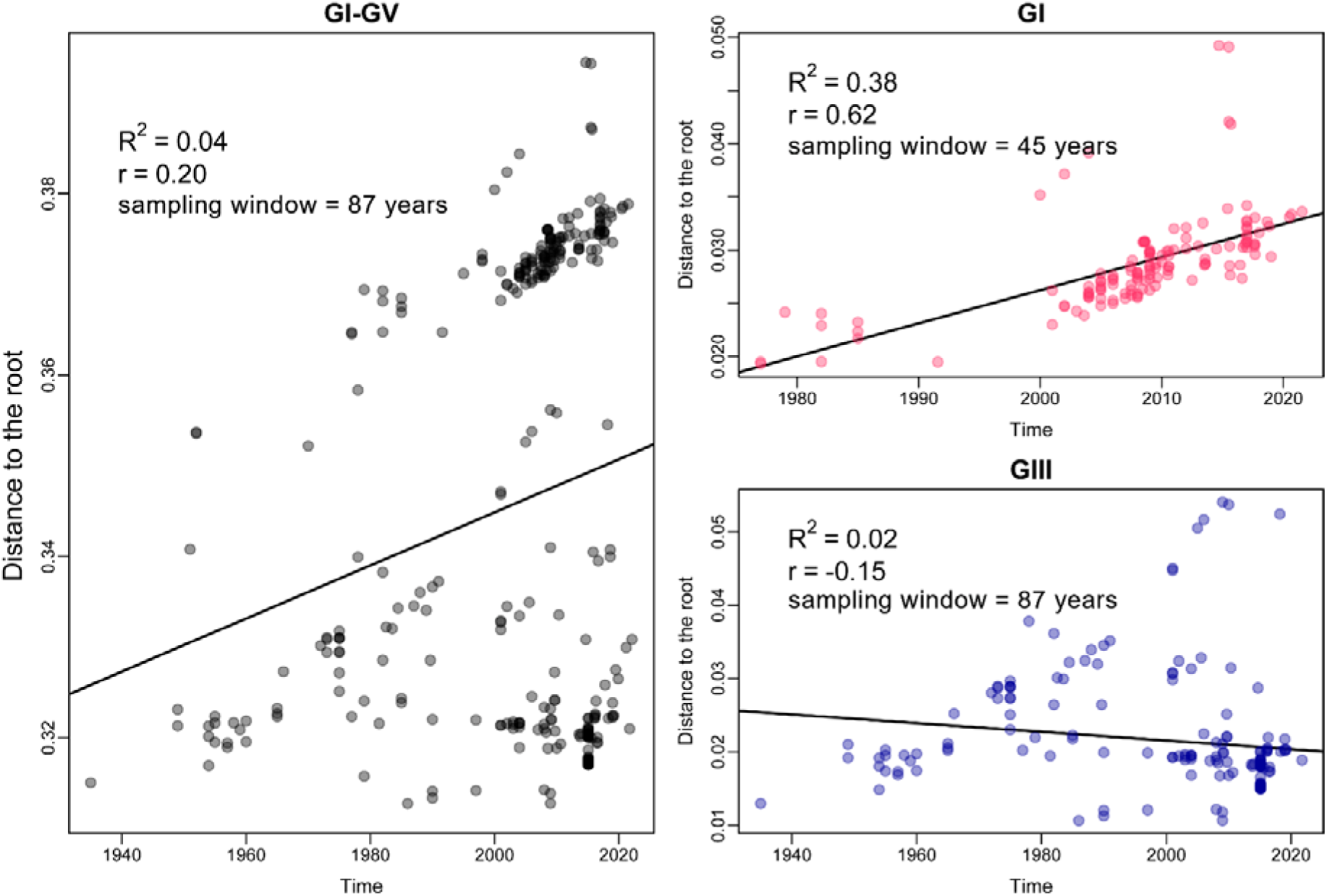
Root-to-tips (RTT) regression analysis of three JEV data sets; X-axes represent calendar years, Y-axes—distances from a root to tips.

For each genotype pair, we randomly sampled mean (μ□, μ□) and standard deviation (σ□, σ□) values from their posterior distributions and generated two sets of 100,000 draws each: *X*_1_∼lognormal(µ_1_, σ_1_) and *X*_2_∼lognormal(*µ*_2_, σ_2_). For each simulated pair (x_1_, *X*_2_), where *X*_1_ E *X*_1_ and *X*_2_ E *X*_2_, we computed the frequency of *X*_1_ > *X*_2_, yielding P(X_1_ > *X*_2_ | data). This procedure was repeated 1000 times to estimate 95% credible intervals (CIs).

## 3. Results

### 3.1 Degree of clocklike behavior of the common JEV data set (GI-GV) and GI and GIII genotypes

In the first step, using three JEV data sets (GI-GV, GI, GIII) we conducted RTT regression to assess to which degree data behave in a clock-like fashion (Figure 2).

The JEV GI-GV and GIII data sets showed R^2^ values close to zero (0.04 and 0.12 with a negative slope, respectively) indicating the weak clocklike behavior of the data which can be explained by the presence of a large amount of rate variation between lineages of the trees (95% HPD of the coefficient of rate variation (CV), 1.04–1.62 for GI-GV and 1.29–2.4 for GIII). GI, on the contrary, demonstrated a stronger temporal structure with R^2^ of 0.79 and the lowest CV values (95% HPD, 0.43–0.68) (Table 3).

**Table 3.** Comparison of the determination coefficients inferred in the root-to-tip regression analysis and the coefficient of variation values.

| Data set | Number of samples | Sampling window, years | Determination coefficient ($R^2$) inferred in RTT analysis <sup>1</sup> | 95% HPD of the coefficient of rate variation <sup>2</sup> | 95% HPD of the substitution rate (s/s/y) <sup>3</sup> |
| --- | --- | --- | --- | --- | --- |
| GI-GV | 325 | 87 | 0.04 | 1.04–1.62 | $1.67 \times 10^{-4}$–$3.14 \times 10^{-4}$ |
| GI | 134 | 45 | 0.79 | 0.43–0.68 | $3.45 \times 10^{-4}$–$4.82 \times 10^{-4}$ |
| GIII | 171 | 87 | 0.12 | 1.29–2.4 | $3.84 \times 10^{-5}$–$8.63 \times 10^{-5}$ |

<sup>1</sup> Root-to-tip (RTT) regression analysis was performed in Clockor2;
<sup>2</sup> 95% HPDs were inferred in BEAST analysis;
<sup>3</sup> S/s/y—substitutions per site per year.

Importantly, the low R^2^ values or a negative slope should not be unequivocally considered as the absence of temporal signal (Rambaut et al. 2016). RTT regression analysis appears to be uninformative when the data were sampled over a relatively narrow sampling window (GI-GV data set) and/or there is large rate variation among lineages (GIII data set) (Duchene et al. 2020). These factors result in decreasing R^2^. Therefore, we applied BETS—a formal test of temporal signal—to assess the statistical support for including sampling times in a Bayesian analysis using both the strict clock and optimized relaxed clock to take into account among-branch rate variation.

### 3.2 Presence of temporal signal in all JEV data sets

BETS displayed very strong support (log Bayes factor > 5) for M_het_ over M_iso_ in each pair of competitive models (Table 4) with the lowest log Bayes factor of 47.8 for the GIII data set (ORC_het_ relative to ORC_iso_) which possessed the largest amount of rate variation among branches (Table 3). Despite the last fact, in the log scale, such log Bayes factor values can be interpreted as overwhelming support for including sequence sampling times into the model (Table 4, Figure 3).

**Figure 3.**
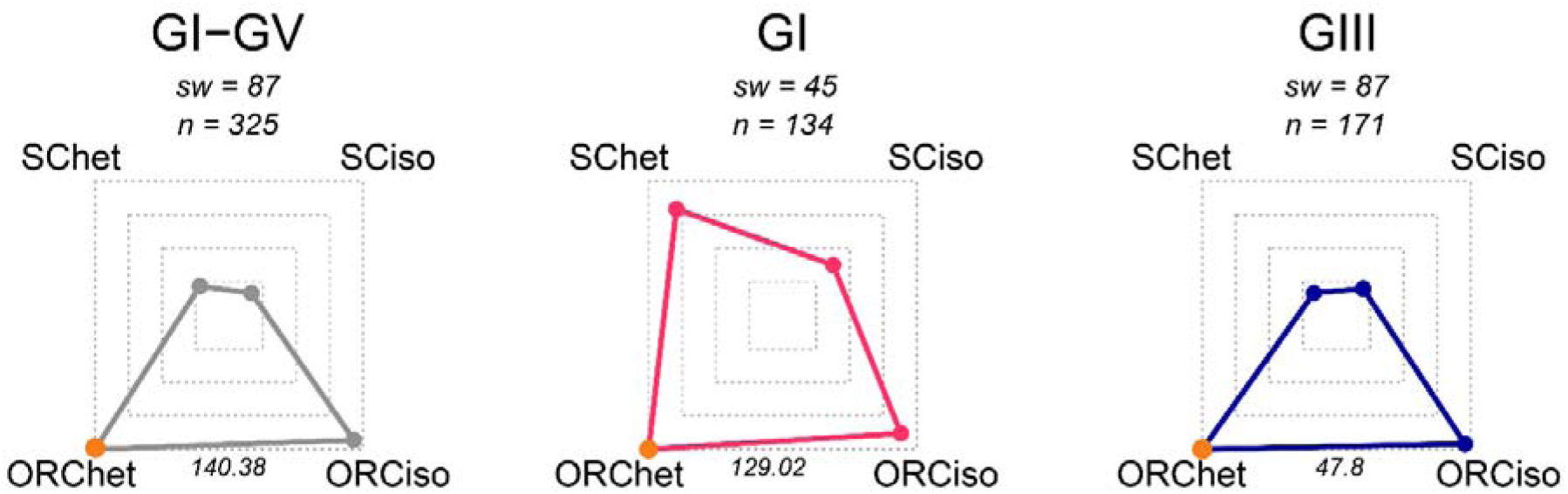
Relative log marginal likelihoods of each JEV data set (GI-GV, GI, GIII) analyzed in our study. Log marginal likelihoods were calculated with four different configurations of sampling times (heterochronous and isochronous trees) and molecular clocks. In the corners, there are models with combinations of molecular clocks and the inclusion or exclusion of sampling times. The outermost dashed borders of the squares correspond to the highest log marginal likelihood calculated. The extent to which the polygons are stretched depicts *relative* model support such that the perfect square indicates equal support for each model. The corner of a polygon that falls close to the center would correspond to a model with the lowest support. SC is a strict clock and ORC is an optimized relaxed clock. *Het* (heterochronous) includes sampling times, while *iso* (isochronous) exclude sampling times. An orange corner points at the model combination with the highest log marginal likelihood. The numbers near the polygon edges are the Bayes factors for the best model (orange corner) over the model with less support. The values above the charts are sampling windows (sw) in years and sample sizes (n).

**Table 4.** Results of the Bayesian test of temporal signal in Japanese encephalitis virus data sets.

| Data set | Molecular clock<sup>1</sup> | Log marginal likelihood | Log Bayes factor<sup>2</sup> |
| --- | --- | --- | --- |
| GI-GV | ORC<sub>iso</sub> | -113559.93 | 140.38 |
|  | ORC<sub>het</sub> | -113419.55 | 0 |
|  | SC<sub>iso</sub> | -115098.41 | 1678.86 |
|  | SC<sub>het</sub> | -114995.46 | 1575.91 |
| GI | ORC<sub>iso</sub> | -54633.51 | 129.02 |
|  | ORC<sub>het</sub> | -54504.49 | 0 |
|  | SC<sub>iso</sub> | -55182.96 | 678.48 |
|  | SC<sub>het</sub> | -54728.94 | 224.45 |
| GIII | ORC<sub>iso</sub> | -46598.66 | 47.80 |
|  | ORC<sub>het</sub> | -46550.86 | 0 |
|  | SC<sub>iso</sub> | -47491.94 | 941.08 |
|  | SC<sub>het</sub> | -47529.78 | 978.92 |

<sup>1</sup> ORC—optimized relaxed clock, SC—strict clock; iso—isochronous model, het—heterochronous model;
<sup>2</sup> Best-performing model has a log Bayes factor value of 0.

The comparison of the SC_het_ and ORC_het_ models also showed very strong evidence in favor of ORC in each data set (Table 4). The lowest log Bayes factor of 224.45 was detected for, as expected, GI which had the highest R^2^ in the RTT regression analysis.

Thus, all three data sets passed the Bayesian test of temporal signal and were analyzed with ORC in the next step of phylogenetic analysis.

### 3.3 Analysis of temporal signal in the previous studies

We analyzed temporal structure in the nine data sets from the previous studies (Figure 4):

**Figure 4.**
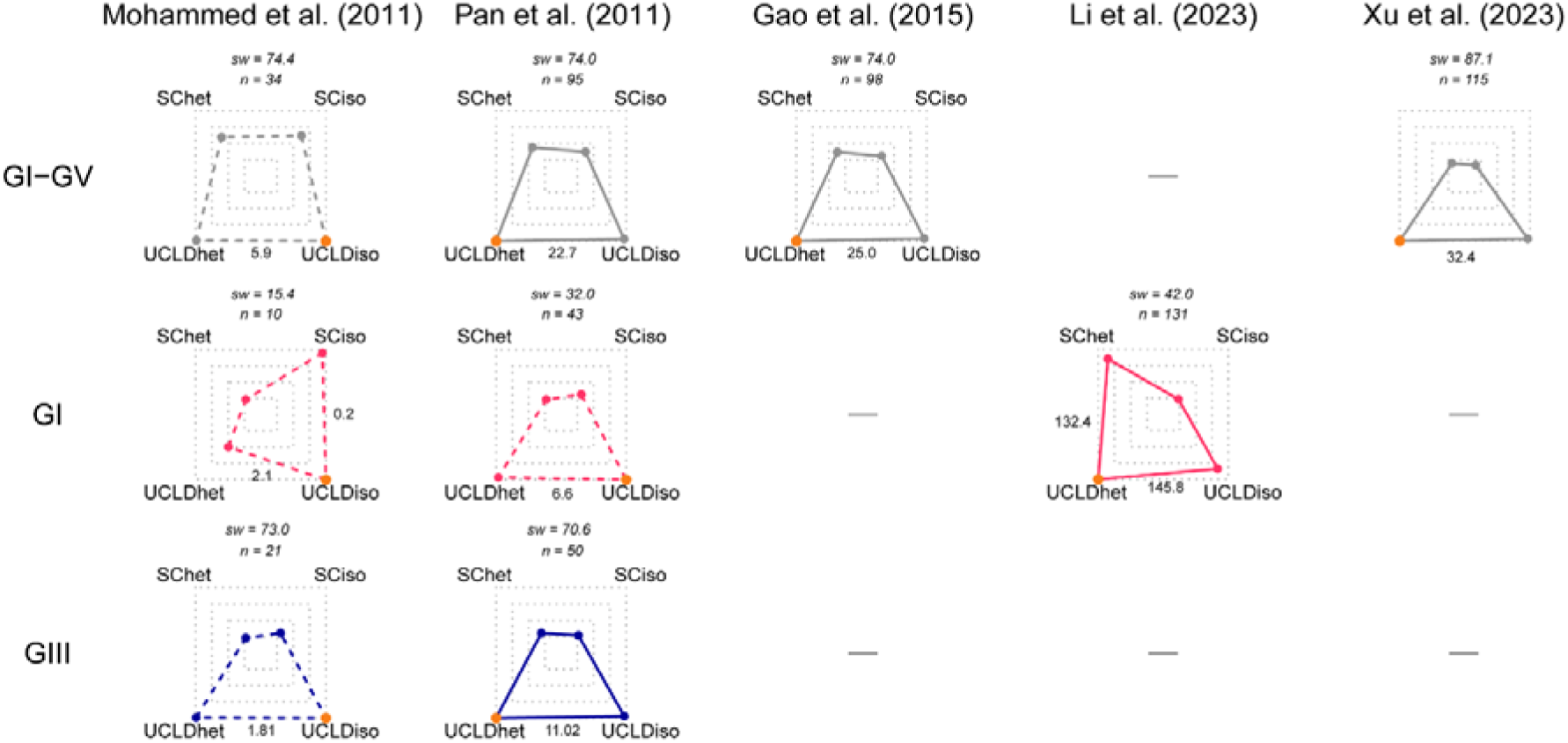
Relative log marginal likelihoods of nine JEV data sets from the previous works (note – data set of a JEV common clade from the study of Pan et al. (2011) exclude the genotype V) calculated with four different configurations of sampling times (heterochronous and isochronous trees) and molecular clocks. The rows of the charts are JEV genotypes, the columns depict the studies from which the data sets were taken. In the corners of each chart, there are models with combinations of molecular clocks and the inclusion or exclusion of sampling times. The extent to which the polygons are stretched depicts *relative* model support such that the perfect square indicates equal support for each model. The outermost dashed borders of the squares correspond to the highest log marginal likelihood. The corner of a polygon that falls close to the center would correspond to a model with the lowest support. SC is a strict clock and UCLD is an uncorrelated log-normal relaxed clock. *Het* (heterochronous) includes sampling times, while *iso* (isochronous) exclude sampling times. An orange corner points at the model combination with the highest log marginal likelihood. The numbers near the polygon edges are the Bayes factors for the best model (orange corner) over the model with less support. The values above the charts are sampling windows (sw) in years and sample sizes (n). The polygons with dashed edges display the data sets with no temporal signal, while solid polygons are the data sets where temporal signal was detected (i.e., heterochronous models including sampling times are favored).

BETS revealed that temporal signal is present in the five of nine JEV data sets (Pan et al. (2011), GI-GIV, GIII; Gao et al. (2015), GI-GV; Li et al. (2023), GI; Xu et al. (2023), GI-GV) with very strong support (the lowest log Bayes factor is 11.02 for JEV GIII from Pan et al. (2011) that contains the lowest number of complete ORF sequences equal to 50). Interestingly, the Bayes factors increased along with sample size (Table S2, Figure S1) which was most likely due to the increase in the number of alignment informative sites. In all data sets from the study of Mohammed et al. (2011) temporal signal was not detected even in the GI-GV data set with 34 complete ORF sequences (log Bayes factor of 5.9 for UCLD_iso_ over UCLD_het_). However, in the case of the GI (n=10) and GIII (n=21) data sets from the same work, BETS displayed only positive support for UCLD_iso_ over UCLD_het_ with the log Bayes factors of 1.8 and 2.1, respectively. The last data set with no temporal signal was JEV GI from the study of Pan et al. (2011) despite the sample size of 43 complete ORF sequences: path sampling showed very strong support for UCLD_iso_ over UCLD_het_ (log Bayes factor = 6.6).

### 3.4 Substitution rates

The mean substitution rate of the common JEV GI-GV clade was 2.41×10^−4^ s/s/y (95% HPD, 1.67×10^−4^–3.14×10^−4^ s/s/y) or approximately 2.5 nucleotide substitutions per ORF (10,296 nucleotides) per year. GI has the highest mean rate of evolution: 4.13×10^−4^ s/s/y (95% HPD, 3.45×10^−4^–4.82×10^−4^ s/s/y) or approximately 4.2 nucleotide substitutions per ORF per year. The lowest mean substitution rate was obtained for GIII: 6.17×10^−5^ s/s/y (95% HPD, 3.84×10^−5^– 8.63×10^−5^ s/s/y) or approximately 0.6 nucleotide substitutions per ORF per year (Figure 5).

**Figure 5.**
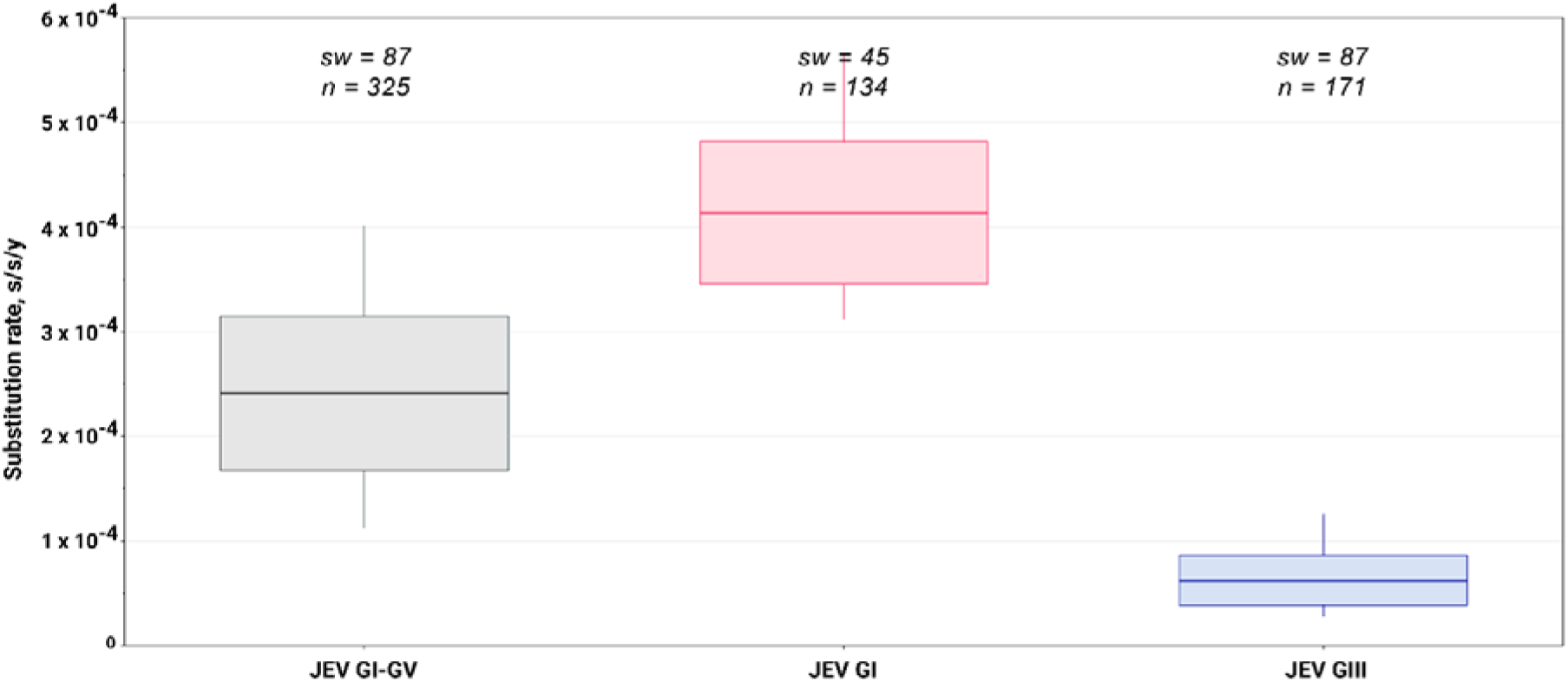
Substitution rates of common JEV clade (GI-GV), genotypes I (GI) and III (GIII). Upper and lower ends of the boxes are 95 % highest posterior density interval borders. The Y-axis represents substitution rate and is expressed in the number of substitutions per site per year (s/s/y). The values above the boxplots are sampling windows (sw) in years and sample sizes (n)

Despite the largest CV (Table 3), the 95% HPD interval for JEV GIII was the narrowest one (Figure 5). The issue of a high-degree of among-branch rate variation in the JEV GIII data set is likely due to passaging some virus strains. In particular, 34 of 171 JEV GIII were isolated from before 1980. In addition, there are clusters with closely related sequences isolated contemporaneously (phylo-temporal structure, Figure 6). Inspecting a rate variation pattern on the MCC tree (Figure S2) revealed the most outstanding substitution rate value of 0.0014 s/s/y on the external branch of BJ-1_BP28 strain (KU871371.1) collected in 2015 (the study has not been published). However, the reanalysis without this strain inferred the nearly identical estimate of mean substitution rate (95% HPD, 3.71×10^−5^– 8.22×10^−5^ s/s/y) (Supplementary Material).

**Figure 6.**
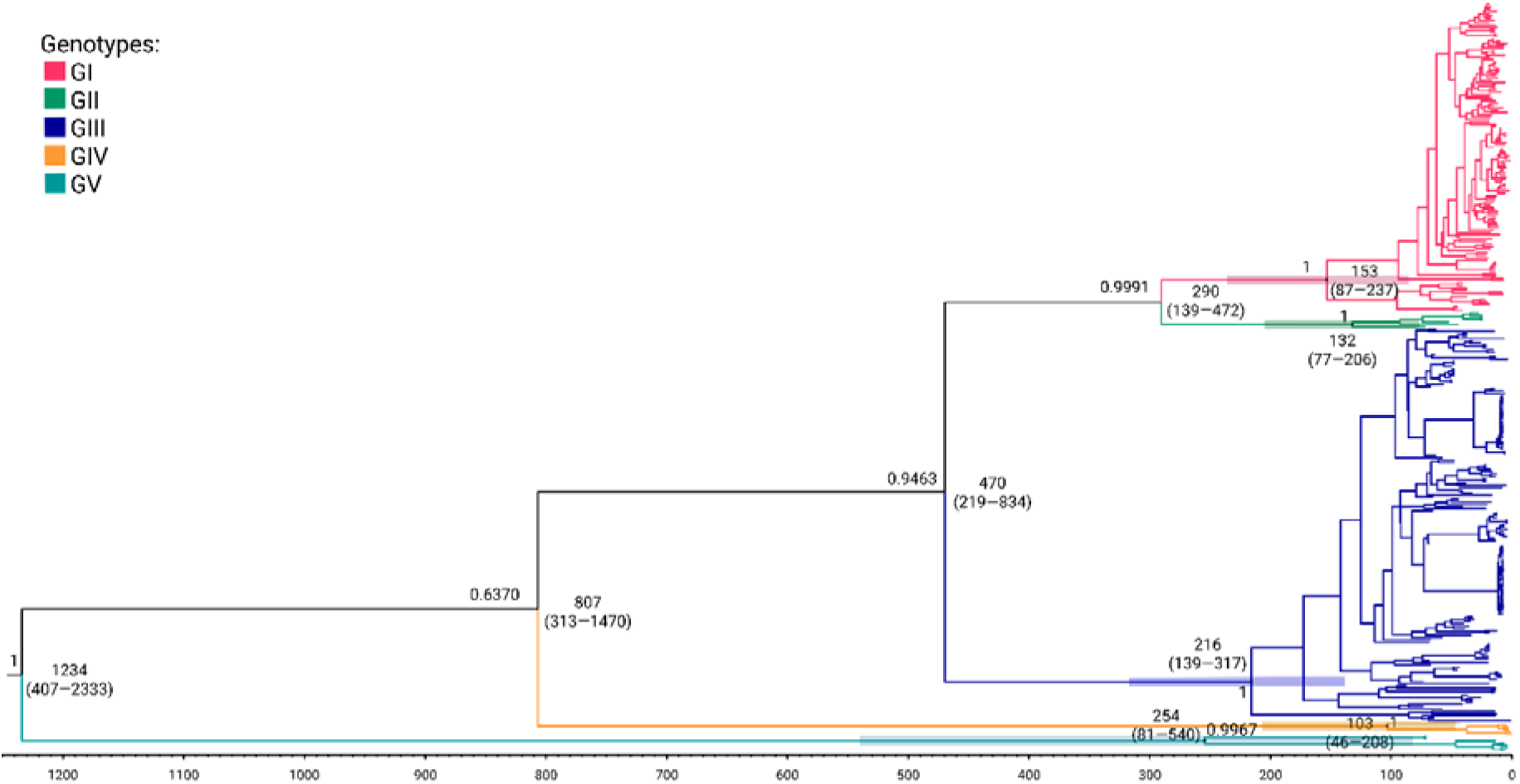
Maximum clade credibility tree of JEV summarized from a posterior tree distribution produced by an MCMC run in BEAST. The numbers nearby the main nodes indicate divergence times before the most recent sample collected on February 20, 2022, with 95% highest posterior density (HPD) intervals within parentheses. The horizontal color bars depict 95% HPD of the divergence times of JEV genotypes.

The mean substitution rate values of the JEV GI-GV clade lies between estimates for JEV GI and GIII and represents the average of rates across branches of the entire tree.

While the 95% HPD intervals of the mean rates do not overlap, among-branch rate variation should be taken into account—particularly for the two data sets (JEV GI-GV and GIII), which exhibit coefficients of rate variation above 1.0 and even 2.0 (Table 3). In other words, some branches of the JEV GIII phylogeny can *probably* exhibit higher rates than those of JEV GI, even though the mean of GIII substantially lower. To resolve this issue, we used 1,000 posterior-sampled mean and sigma values to perform Monte Carlo simulations in R, calculating the posterior probability that a rate drawn from a lognormal distribution of one genotype would exceed that of another genotype. We tested the following three pairs of JEV data sets (Table 5):

**Table 5.** Posterior probability that a lineage from Genotype 1 evolves faster than that of Genotype 2 approximated by Monte Carlo simulations in R.

| Genotype 1 ($X_1$) | Genotype 2 ($X_2$) | 95% CI for $P(X_1 > X_2 \mid \text{data})^*$ |
| --- | --- | --- |
| JEV GI | JEV GIII | 0.64–0.79 |
| JEV GI | JEV GI-GV | 0.87–0.97 |
| JEV GI-GV | JEV GIII | 0.69–0.84 |

The results of the comparison indicate that while JEV GI, on average, evolves faster than JEV GIII and the common JEV GI-GV clade, the reverse scenario can also occur with some probability. In the pair of JEV GI and JEV GIII, the 95% CI (0.64–0.79) indicates a 21–36% probability that rates from the lognormal distribution of JEV GIII exceed those from JEV GI although the mean evolutionary rate of JEV GI is substantially higher than that of JEV GIII (Figure 5). This is probably due to the largest coefficient of variation of JEV GIII (1.29– 2.4). The similar observation was made for the comparison of evolutionary rates of JEV GI-GV > JEV GIII (0.69–0.84).

For the JEV GI versus JEV GI-GV comparison, the 95% credible interval (0.87–0.97) indicates a high probability that a rate sampled from the JEV GI distribution would exceed that of the JEV GI-GV common clade distribution.

### 3.5 Divergence times

The evaluation of JEV divergence times was performed using 325 complete ORF nucleotide sequences (Figure 6). The results of the analysis in BEAST suggest that the tMRCA of JEV is 1234 years before 2022 (95% HPD, 407–2333 years).

Concerning the emergence of JEV genotypes, 95% HPD intervals of time to MRCA of all JEV genotypes are overlapped (Figure 6). Therefore, we are unable to reliably establish the chronology of the occurrence of genotypes. Besides, only the GI and GIII are representative in terms of sample size, and, as a consequence, the estimates of the other genotype ages (GII, GIV, GV) are unreliable. GI and GIII ages, in their turn, are not statistically indistinguishable: GI occurred 153 years ago (95% HPD, 87–237) and GIII—216 years ago (95% HPD, 139–317). Generally, considering the bounds of 95 HPDs obtained, JEV genotypes occurred 77–540 years ago.

## 4. Discussion

Our study demonstrates the first assessment of JEV evolutionary rate supported by the formal analysis of temporal signal in the nucleotide data. In the literature, there is substantial discrepancy between JEV substitution rate values estimated by different authors. Therefore, we also evaluated temporal structure in the data sets from the previously reported studies. Our findings indicate which of those estimates are reliable and which were based on uninformative data and/or on the priors set in a program by the researchers on their own. The results of our analysis show that JEV GI is the fastest-evolving genotype and, conversely, GIII is the slowest JEV genotype. Regarding the width of 95% HPD interval, our JEV rate estimates are among the most precise.

### 4.1 Comparing the mean substitution rates of JEV yielded in our analysis with estimates reported previously

We compared the JEV substitution rate values obtained in our study with the previous estimates based on the analysis of complete ORF gene sequences (Figure 7).

**Figure 7.**
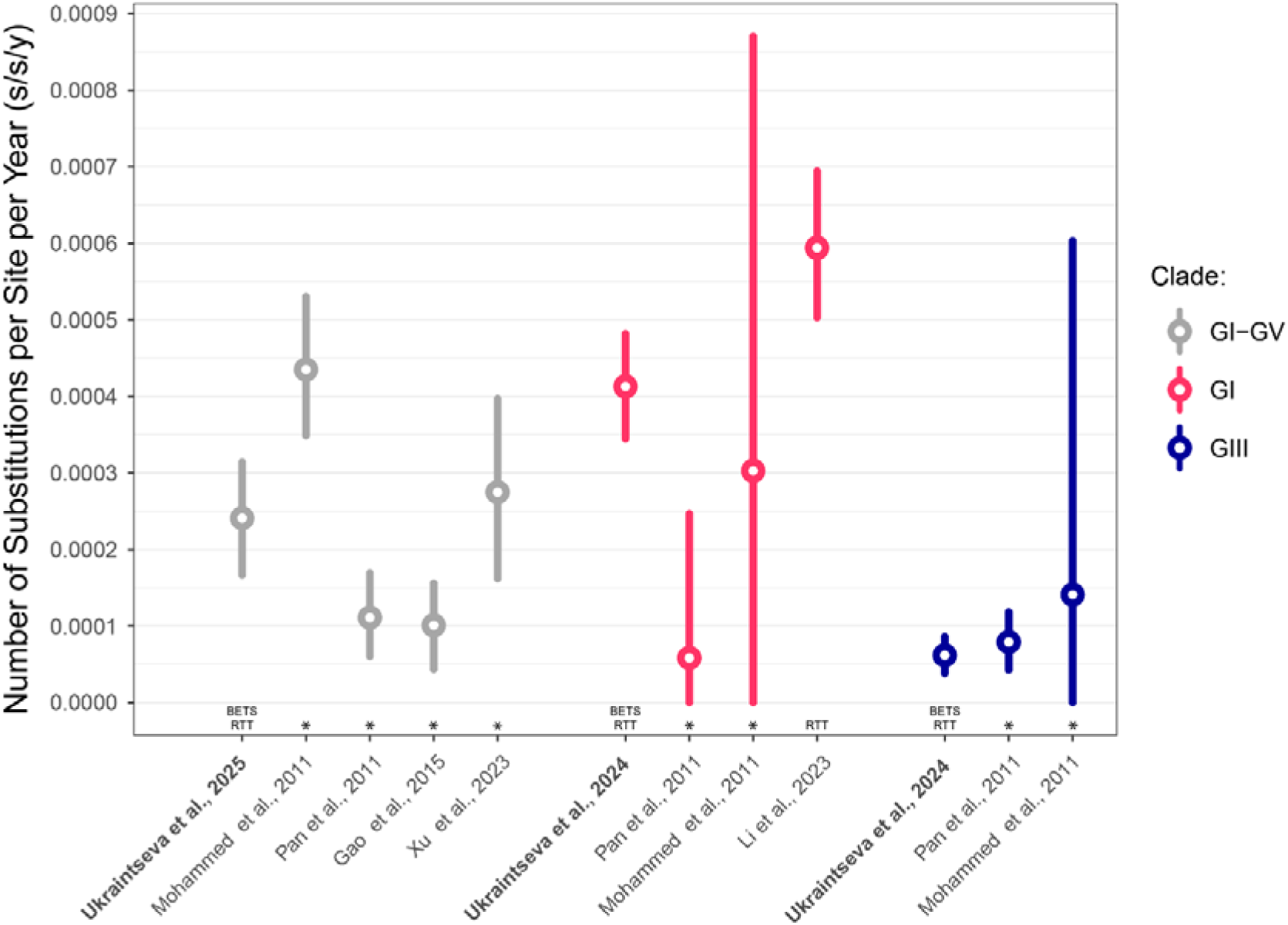
Comparison of the Japanese encephalitis virus (JEV) substitution rate evaluated in our study based on complete open reading frame nucleotide sequences and the JEV substitution rates according to the literature. The vertical bars represent 95% highest posterior density intervals, circles—median or mean values. The X-axis labels are references to appropriate studies, the methods on temporal signal evaluation applied are indicated above the X-axis: BETS—Bayesian evaluation of temporal signal, RTT—root-to-tips regression, *— temporal signal was not assessed.

First of all, we found only one study where at least the degree of clock-like behavior was assessed with informal RTT (Li et al. 2023) when analyzing GI. Therefore, the substitution rate values reported in the literature should be treated *with caution*, especially in the cases where the number of samples was relatively low. Hence, to our knowledge, we provide here, for the first time, the JEV evolutionary rate estimates confirmed by a formal statistical test. Besides, we additionally assessed temporal signal with BETS in the data sets from the previous studies to improve the rigor of the manuscript and attempt to explain the difference between rate estimates observed. Four of nine data sets analyzed displayed the absence of temporal signal (Figure 4, Table S2). Our results emphasize the importance of temporal signal evaluation in the data sets represented by dozens of sequences, even those that are complete or near complete genomes.

#### 4.1.1 JEV GI-GV clade

The comparison showed that the evolutionary rate of the JEV common clade including all five genotypes GI-GV obtained in our study (95% HPD, 1.67×10^−4^– 3.14×10^−4^ s/s/y) is substantially lower than values reported by Mohammed et al. (2011) (95% HPD, 3.49×10^−4^–5.30×10^−4^) who analyzed the data set with the smaller sample size (n=35). Our evaluation of temporal signal in the data set of JEV GI-GV from the study of Mohammed et al. (2011) revealed the absence of temporal signal with very strong support for the isochronous models over heterochronous ones (log Bayes factor = 6.6) (Figure 4) which explained the discrepancy between estimates inferred in our work and in the study of Mohammed et al. (2011).The fact is that, in the case of low-informative and uninformative data, the Bayesian analysis turns out to be very sensitive to a prior choice. In particular, under the coalescent process an inferred substitution rate is largely rested on, for example, a continuous-time Markov chain (CTMC) rate reference prior (default prior on the clock rate in BEAST1). The CTMC prior is a gamma probability distribution with a shape parameter α=0.5 and rate parameter β=Τ, where Τ is tree length, meaning that the CTMC is ultimately conditioned by an effective population size prior (under the coalescent process the expected time of divergence between lineages is inversely proportional to the population size) (Ferreira and Suchard 2010). In brief, the more population size, the longer tree and, as a consequence, the slower evolutionary rate estimated. Thus, interactions between the population size and CTMC priors may drastically affect the inference of substitution rates (we demonstrated this effect using the data set of JEV GI from the study of Mohammed et al. (2011), Figure S3).

In contrast, our estimates are substantially higher than the JEV substitution rates provided by Pan et al. (2011) (95% HPD, 6.04×10^−5^–1.69×10^−4^) and Gao et al. (2015) (95% HPD, 4.37×10^−5^–1.56×10^−4^) who analyzed 98 (GI-GIV) and 100 (GI-GV) sequences, respectively. Our analysis of temporal signal in the data sets from those works revealed that BETS supported the UCLD model with sampling times (log Bayes factors of 22.7 and 25.0 for UCLD_het_ over UCLD_iso_, respectively) (Figure 4). The discrepancy in the substitution rate evaluation between our and the previous works can be explained by the difference in data sets and sample sizes. This is particularly important in the event of high degree of among-lineage rate variation since excluding or including certain tips can drastically affect inferred CV values and, as a consequence, virus evolutionary rate. In the case of the study of Pan et al. (2011), the authors did not include JEV GV in the analysis of the common clade of JEV which also explain the discrepancy between the estimates of a JEV substitution rate.

In one of the last studies, Xu et al. (2023) reported a JEV GI-GV substitution rate (95% HPD, 1.62×10^−4^–3.97×10^−4^) comparable with our estimates. BETS also demonstrated strong support for temporal signal in this data set (Figure 4) with the log Bayes factor of 32.4 for UCLD_het_ over UCLD_iso_. Notably, the sample size (115), clock model (UCLD) and tree prior (Bayesian Skyline) applied in the work of Xu et al. (2023) were similar to two previously discussed studies with substantially lower JEV substitution rates (Pan et al. 2011; Gao et al. 2015). The possible explanation is that Xu et al. (2023) provide new JEV nucleotide sequences, which were not used in our study and in the previous works of the other authors.

#### 4.1.2 JEV GI clade

Our estimates of JEV GI evolutionary rate (95% HPD, 3.45×10^−4^– 4.82×10^−4^) are substantially higher than those we calculated for the GI-GV clade (95% HPD, 1.67×10^−4^–3.14×10^−4^) (Figure 5). JEV GI rate values inferred in our study are also substantially different from the literature data, except for the estimates with the widest 95% HPD interval (2.09×10^−8^ 8.7×10^−4^) provided by Mohammed et al. (2011) using only 10 complete ORF sequences. BETS revealed the absence of temporal structure in this data (Figure 4) with log Bayes factor for UCLD_iso_ over UCLD_het_ of 2.1 (positive evidence).

The study of Pan et al. (2011) (n=43) reported the substitution rate estimates with an extremely low border of 95% HPD (3.42^−9^–2.47×10^−4^), which indicates weak or absent temporal signal. In our evaluation of temporal structure in this data set, the UCLD model in which sampling times are contemporaneous (UCLD_iso_) was favored with very strong support (log Bayes factor = 6.6) over the UCLD model in which the data was accompanied by the actual sampling times (UCLD_het_) (Figure 4).

The only study (Li et al. 2023) in which the extent of time-dependency in data was assessed (RTT regression analysis) reported statistically higher JEV GI evolutionary rates (95% HPD, 5.03×10^−4^–6.94×10^−4^) than our estimates. Our assessment of temporal signal in this data set showed very strong support for the presence of temporal signal (log Bayes factor for UCLD_het_ over UCLD_iso_ of 145.8). We speculated on three possible explanations of the discrepancy—the first one is the authors removed RTT regression outliers without explaining the reason (e.g., sequencing errors, long passage history, lab contamination, etc.) that is forcing the data to fit a strict clock model, which does not assume among-lineage rate variation, instead of using an alternative model (e.g., relaxed clock) to describe the data; the second one is the new JEV nucleotide sequences generated in the study of Li et al. (2023) and not included in our work; the third reason is the smaller sample size (n=131) compared to our work.

#### 4.1.3 JEV GIII clade

Among all three data sets analyzed (Table 3), JEV GIII was characterized by the lowest mean evolutionary rate (95% HPD, 3.84×10^−5^–8.63×10^−5^) (Figure 4). Thus, JEV GIII evolves generally on the other order of magnitude than JEV GI and JEV GI-GV.

The very similar estimates (4.28×10^−5^–1.18×10^−4^) were reported by Pan et al. (2011) with a sample size of 53 and a model combination comparable to the one we applied in our work with the exception of a relaxed molecular clock (UCLD). Our analysis of temporal structure with BETS very strongly supported heterochronous model over an isochronous one (log Bayes factor of 11.02). In the study of Mohammed et al. (2011), the 95% HPD interval was the widest one (7.05×10^−8^–6.03×10^−4^). As in the case of JEV GI, BETS indicated the absence of temporal signal in the JEV GIII data set from the work of Mohammed et al. (2011) and the main reason is the small sample size (n=22) and, therefore, the small number of variable sites in the alignment.

However, it is worth noting that regardless of precise estimates, the JEV GIII data set includes a large number of strains isolated before the 1980s, which could be serially passaged over a period of decades before being sequenced. This leads to artificially accelerated accumulation nucleotide substitutions and, as a result, to the overestimation of a substitution rate. Also, the long passage history is the most likely cause of the high CV values (Table 3) and the RTT negative slope (Figure 2) inferred. Thus, the true JEV GIII evolutionary rate is likely *lower* than the values reported in our study.

### 4.2 Comparing the substitution rates of JEV (GI-GV clade) and other mosquito-borne flaviviruses

To provide a better understanding of the JEV evolutionary rate, we compared the rate of JEV GI-GV clade obtained with rates of JEV’s most closely related species and the other well-studied MBFV (Figure 8).

**Figure 8.**
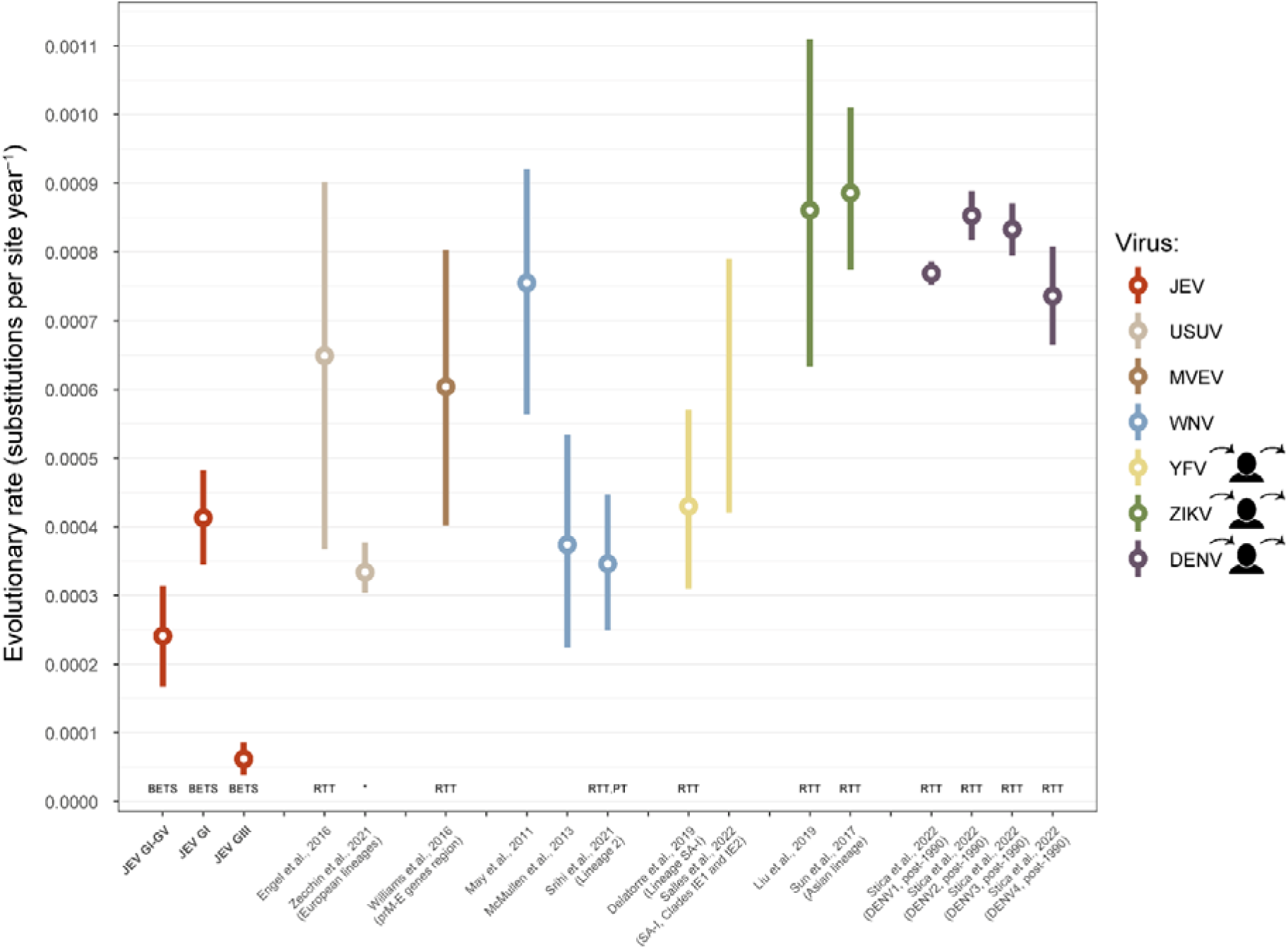
Comparison of the Japanese encephalitis virus (JEV) substitution rate inferred in our study using complete open reading frame nucleotide sequences of GI–GV, GI, and GIII genotypes, and the substitution rates of the other mosquito-borne flaviviruses according to the literature: USUV—Usutu virus (Engel et al. 2016; Zecchin et al. 2021), MVEV—Murray Valley encephalitis virus (Williams et al. 2015), WNV—West Nile virus (May et al. 2011; McMullen et al. 2013; Srihi et al. 2021), YFV—yellow fever virus (Delatorre et al. 2019; Moreira Salles et al. 2022), ZIKV—Zika virus (Sun et al. 2017; Liu et al. 2019), DENV—dengue virus (Stica et al. 2022). The person icons to the right of the abbreviations point to the viruses for which humans are not dead-end hosts. The vertical bars represent 95% highest posterior density intervals, circles—median or mean values if available. The X-axis labels are references to appropriate studies. The methods on temporal signal evaluation applied are indicated above the X-axis: BETS—Bayesian evaluation of temporal signal, RTT—root-to-tips regression, PT—permutation test, *—temporal signal was not assessed.

To compare the JEV substitution rate, we considered the data only if 95% HPD intervals for mean/median were provided. Also, the estimates reported with mistakes or obvious typos were excluded from the comparison. We mainly focused on the substitution rates derived from the analysis of as many virus genotypes as possible (i.e., the genetic diversity of viruses was meant to be represented to its greatest extent).

Unlike the literature data on JEV, the most of MBFV rate estimates were supported by the RTT regressions, and, furthermore, a permutation or randomization test was applied in one study (Srihi et al. 2021). This makes the comparison much more reliable, but, for some virus species, reported rates are statistically different between studies.

First, we compared our JEV substitution rate estimates with the rates of JEV most closely related species (Moureau et al. 2015)—Usutu virus (USUV), Murray Valley encephalitis virus (MVEV), West Nile virus (WNV)—for which the sample size allows the evolutionary rate of the virus to be reliably evaluated. Second, we included in the comparison the most widely understood members of MBFVs— yellow fever virus (YFV), Zika virus (ZIKV), and dengue virus (DENV).

As shown in Figure 8, JEV is one of the slowest changing MBFVs represented in our analysis (95% HPD, 1.67×10^−4^–3.14×10^−4^). However, when accounting for among-branch rate variation (Tables 3 and 5), JEV can produce branches with evolutionary rates that exceed the mean rate of JEV GI, bringing it closer to the other MBFVs.

The overlap of 95% HPD intervals is observed in the case of two assessments of WNV (2.24×10^−4^–5.34×10^−4^ according to McMullen et al. (2013) and 2.50×10^−4^–4.47×10^−4^ for WNV lineage 2 reported by Srihi et al. (2021)). There is a slight overlap with the rate values of the USUV European lineage (95% HPD, 3.04×10^−4^–3.77×10^−4^). More distant MBFV members evolve faster, especially ZIKV (Sun et al. 2017; Liu et al. 2019) and DENV (Stica et al. 2022), the substitution rates of which are near the threshold of 10^−3^ and 10^−2^ orders of magnitude or even above it.

The low substitution rate of virus can play a positive role in the virus strain selection for the vaccine production. Vaccines for slowly evolving viruses remain effective longer. To date, there are four types of effective JEV vaccines based on the GIII strain (Srivastava et al. 2023). Regarding the more rapidly changing viruses, as a rule, there is either no effective vaccine (e.g., ZIKV), or in the case of DENV the vaccinated person had to be previously infected with the virus (https://www.who.int/publications/i/item/who-wer-9918-203-224) (although there is an effective vaccine against poliovirus despite the virus exhibiting a high evolutionary rate (Guo et al. 2023)). Likewise, the vaccine strain should be constantly updated as in the case of SARS-CoV-2 or influenza viruses.

Rates of evolution are determined by many factors including mutation rate (influenced, for instance, by RNA replication errors), natural selection, generation time (time between successive infections in transmission chains) and population size (Duchene et al. 2016). There is also a negative correlation between virus genome size and evolutionary rate (Ho 2020). Considering RNA viruses which exhibit evolutionary rate between 10^−4^ and 10^−3^ orders of magnitude (Duchene et al. 2014), the mean substitution rate of JEV (95% HPD, 1.67×10^−4^–3.14×10^−4^ s/s/y) is relatively slow. The reason why the JEV evolution rate was slower than its closest relatives is unclear and the answer likely lies in the complex interaction of the described factors. We assume that humans which are not dead-end hosts (Figure 8) in the case of fastest evolving viruses (YFV, ZIK, DENV) may shorten the time between successive infections. However, USUV and MVEV viruses demonstrate relatively high evolutionary rates, (although, calculated without a formal test of temporal signal). Therefore, humans as amplifying hosts can play a rather indirect role in virus viability.

### 4.3 Divergence time

Despite the relatively large number of complete ORF sequences used in our study, the uncertainty in the assessment of the evolutionary timescale of the entire tree spans a large time interval (95% HPD of root height, 407–2333 years). A little more precise estimates of time to the most recent common ancestor (MRCA) of JEV GI-GV (95% HPD, 463–2100 years) were reported by Xu et al. (2023) with a sample size of 115 complete genome sequences. In the other studies with smaller sample sizes, 95% HPD intervals for tree roots were much wider: Gao et al. (2015) conclude that the MRCA of JEV GI-GV is estimated to have occurred 3255 years ago (95% HPD, 978–6125 years) encompassing much of the range of our estimate; Pan et al. (2011) studied JEV GI-GIV defined the MRCA age of the entire clade as 1695 years ago (95% HPD, 548–3153 years). More recent JEV emergence was described by Mohammed et al. (2011) (95% HPD, 374–500 years ago), but the sample size was the smallest one (n=35) which implies lack of temporal structure in the data and is also reflected in the substitution rate estimates (Figure 7).

The yielded ages of JEV subtypes are consistent with the previous assessments (Xu et al. 2023). However, the “minor” JEV genotypes, such as GII, GIV, GV, were mainly scrutinized in the studies of E gene data sets. The estimates of time to MRCA based on E gene sequences are more precise (Schuh et al. 2013; Sikazwe et al. 2022), whereas the substitution rate of E gene is substantially higher (95% HPD, 3.92×10^−4^–6.52×10^−4^) and MRCA ages are younger (e.g., the MRCA of JEV GI-GV, 95% HPD 216–921 years) (Schuh et al. 2013); JEV GI, 44–55 years (Li et al. 2025)). Rejuvenation of the tree root along with (or possibly due to) an increase in the evolutionary rate indicates the impact of the priors on the inference of the evolutionary timescale. In other words, even if the E gene of JEV evolved faster than the entire ORF gene, then the dates of the emergence of JEV MRCA should not substantially differ between the analyses of two loci in the case of the obligatory presence of temporal signal in both data sets. However, the sample sizes also can play a significant role. In the cited works dedicated to the E gene analysis, only Sikazwe et al. (2022) performed the RTT analysis, however, JEV GV samples were removed from the analysis as outliers without a justified reason. Moreover, the R^2^ and the regression visualization were not reported as well as the substitution rate of the common data set (JEV GI-GIV). Thus, the problem of the fitness of the E gene for JEV date-stamped phylogeny reconstruction should be addressed in future studies.

## Conclusions

1. All complete ORF data sets analyzed (JEV GI-GV, GI, GIII) yielded strong temporal signal in BETS. It means that a sufficient amount of molecular evolution occurred over the sampling time window to permit collection dates of sequences for clock calibration. Thus, JEV population can be treated as measurably evolving and the reconstructions of JEV dated phylogenies is warranted and reliable;
2. JEV represented by the clade comprising all five genotypes (GI-GV) has the lowest mean substitution rate in comparison to its most closely related species and other well-studied members of MBFV: 2.41×10^−4^ s/s/y (95% HPD, 1.67×10^−4^–3.14×10^−4^ s/s/y) or approximately 2.5 nucleotide substitutions per ORF (10,296 nucleotides) per year. To our knowledge, it is the first assessment of the JEV substitution rate confirmed by the formal Bayesian test of temporal signal;
3. JEV GIII is characterized by the lowest mean substitution rate (6.17×10^−5^ s/s/y (95% HPD, 3.84×10^−5^–8.63×10^−5^ s/s/y) or approximately 0.6 nucleotide substitutions per ORF per year) in comparison with GI and the common JEV clade. However, the passage history of a large number of strains was not taken into account, hence the true evolutionary rate of JEV GIII. Furthermore, among-lineage rate variation also plays a significant role. The Monte Carlo simulations showed a 21–36% posterior probability that GIII substitution rates would exceed those of GI despite GI having the highest mean;
4. Time to JEV MRCA (GI-GV clade) has the mean value of 1234 years (95% HPD, 407–2333 years);
5. The mean values and appropriate 95% HPD intervals of divergence of each JEV genotype are comparable and close. Based on the lower and upper bounds of HPDs, JEV genotypes occurred in the period from 77 to 540 years ago. The main limitations of our study are that JEV GII, GIV, and GV genotypes represent an insufficient number of complete or near complete genomes. Therefore, future studies including more JEV genomes will obtain more reliable and precise substitution rate estimates given that temporal signal will access in proper way.

## Supporting information

https://doi.org/10.6084/m9.figshare.25486117

## References

Ayres DL, Darling A, Zwickl DJ, Beerli P, Holder MT, Lewis PO, Huelsenbeck JP, Ronquist F, Swofford DL, Cummings MP, Rambaut A, Suchard MA (2012) BEAGLE: an application programming interface and high-performance computing library for statistical phylogenetics. Syst Biol 61:170–173. 10.1093/sysbio/syr100

Bouckaert R, Vaughan TG, Barido-Sottani J, Duchene S, Fourment M, Gavryushkina A, Heled J, Jones G, Kuhnert D, De Maio N, Matschiner M, Mendes FK, Muller NF, Ogilvie HA, du Plessis L, Popinga A, Rambaut A, Rasmussen D, Siveroni I, Suchard MA, Wu CH, Xie D, Zhang C, Stadler T, Drummond AJ (2019) BEAST 2.5: An advanced software platform for Bayesian evolutionary analysis. PLoS Comput. Biol 15:e1006650. 10.1371/journal.pcbi.1006650

Crits-Christoph A, Levy JI, Pekar JE, Goldstein SA, Singh R, Hensel Z, Gangavarapu K, Rogers MB, Moshiri N, Garry RF, Holmes EC, Koopmans MPG, Lemey P, Peacock TP, Popescu S, Rambaut A, Robertson DL, Suchard MA, Wertheim JO, Rasmussen AL, Andersen KG, Worobey M, Debarre F (2024) Genetic tracing of market wildlife and viruses at the epicenter of the COVID-19 pandemic. Cell 187:5468–5482 e5411. 10.1016/j.cell.2024.08.010

Delatorre E, de Abreu FVS, Ribeiro IP, Gomez MM, Dos Santos AAC, Ferreira-de-Brito A, Neves M, Bonelly I, de Miranda RM, Furtado ND, Raphael LMS, da Silva LFF, de Castro MG, Ramos DG, Romano APM, Kallas EG, Vicente ACP, Bello G, Lourenco-de-Oliveira R, Bonaldo MC (2019) Distinct YFV Lineages Co-circulated in the Central-Western and Southeastern Brazilian Regions From 2015 to 2018. Front Microbiol 10:1079. 10.3389/fmicb.2019.01079

Douglas J, Zhang R, Bouckaert R (2021) Adaptive dating and fast proposals: Revisiting the phylogenetic relaxed clock model. PLoS Comput Biol 17:e1008322. 10.1371/journal.pcbi.1008322

Duchene S, Holmes EC, Ho SY (2014) Analyses of evolutionary dynamics in viruses are hindered by a time-dependent bias in rate estimates. Proc Biol Sci 281. 10.1098/rspb.2014.0732

Duchene S, Holt KE, Weill FX, Le Hello S, Hawkey J, Edwards DJ, Fourment M, Holmes EC (2016) Genome-scale rates of evolutionary change in bacteria. Microb Genom 2:e000094. 10.1099/mgen.0.000094

Duchene S, Lemey P, Stadler T, Ho SYW, Duchene DA, Dhanasekaran V, Baele G (2020) Bayesian Evaluation of Temporal Signal in Measurably Evolving Populations. Molecular Biology and Evolution. 10.1093/molbev/msaa163

Duffy S, Shackelton LA, Holmes EC (2008) Rates of evolutionary change in viruses: patterns and determinants. Nat Rev Genet 9:267–276. 10.1038/nrg2323

Engel D, Jost H, Wink M, Borstler J, Bosch S, Garigliany MM, Jost A, Czajka C, Luhken R, Ziegler U, Groschup MH, Pfeffer M, Becker N, Cadar D, Schmidt-Chanasit J (2016) Reconstruction of the Evolutionary History and Dispersal of Usutu Virus, a Neglected Emerging Arbovirus in Europe and Africa. mBio 7:e01938–01915. 10.1128/mBio.01938-15

Featherstone LA, Rambaut A, Duchene S, Wirth W (2024) Clockor2: Inferring global and local strict molecular clocks using root-to-tip regression. Syst Biol. 10.1093/sysbio/syae003

Ferreira MAR, Suchard MA (2010) Bayesian analysis of elapsed times in continuous-time Markov chains. Canadian Journal of Statistics 36:355–368. 10.1002/cjs.5550360302

Gao X, Liu H, Li M, Fu S, Liang G (2015) Insights into the evolutionary history of Japanese encephalitis virus (JEV) based on whole-genome sequences comprising the five genotypes. Virol J 12:43. 10.1186/s12985-015-0270-z

Guo Q, Zhu S, Wang D, Li X, Zhu H, Song Y, Liu X, Xiao F, Zhao H, Lu H, Xiao J, Yu L, Wang W, He Y, Liu Y, Li J, Zhang Y, Xu W, Yan D (2023) Genetic characterization and molecular evolution of type 3 vaccine-derived polioviruses from an immunodeficient patient in China. Virus Res 334:199177. 10.1016/j.virusres.2023.199177

Ho SYW (2020) The Molecular Clock and Evolutionary Rates Across the Tree of Life. In: Ho SYW (ed) The Molecular Evolutionary Clock. Springer Cham

Holmes EC, Dudas G, Rambaut A, Andersen KG (2016) The evolution of Ebola virus: Insights from the 2013-2016 epidemic. Nature 538:193–200. 10.1038/nature19790

Kalyaanamoorthy S, Minh BQ, Wong TKF, von Haeseler A, Jermiin LS (2017) ModelFinder: fast model selection for accurate phylogenetic estimates. Nat. Methods 14:587–589. 10.1038/nmeth.4285

Kass RE, Raftery AE (1995) Bayes Factors. Journal of the American Statistical Association 90:773–795. 10.1080/01621459.1995.10476572

Katoh K, Standley DM (2013) MAFFT multiple sequence alignment software version 7: improvements in performance and usability. Mol Biol Evol 30:772–780. 10.1093/molbev/mst010

Ladreyt H, Durand B, Dussart P, Chevalier V (2019) How Central Is the Domestic Pig in the Epidemiological Cycle of Japanese Encephalitis Virus? A Review of Scientific Evidence and Implications for Disease Control. Viruses 11. 10.3390/v11100949

Larsson A (2014) AliView: a fast and lightweight alignment viewer and editor for large data sets. Bioinformatics 30:3276–3278. 10.1093/bioinformatics/btu531

Li F, Feng Y, Wang G, Zhang W, Fu S, Wang Z, Yin Q, Nie K, Yan J, Deng X, He Y, Liang L, Xu S, Wang Z, Liang G, Wang H (2023) Tracing the spatiotemporal phylodynamics of Japanese encephalitis virus genotype I throughout Asia and the western Pacific. PLoS Negl Trop Dis 17:e0011192. 10.1371/journal.pntd.0011192

Li G, Li X, Chen J, Lemey P, Vrancken B, Su S, Dellicour S, Gámbaro F (2025) Tracing more than two decades of Japanese encephalitis virus circulation in mainland China. Journal of Virology 99:e01575–01524. doi:10.1128/jvi.01575-24

Liu RF, He ZJ, Mei P, Xi JC, Cao XD, Yuan LH, Lu JH (2019) Risk assessment and genomic characterization of Zika virus in China and its surrounding areas. Chin Med J (Engl) 132:1645–1653. 10.1097/CM9.0000000000000317

Mackenzie JS (2005) Emerging zoonotic encephalitis viruses: lessons from Southeast Asia and Oceania. J Neurovirol 11:434–440. 10.1080/13550280591002487

Martin DP, Varsani A, Roumagnac P, Botha G, Maslamoney S, Schwab T, Kelz Z, Kumar V, Murrell B (2021) RDP5: a computer program for analyzing recombination in, and removing signals of recombination from, nucleotide sequence datasets. Virus Evol 7:veaa087. 10.1093/ve/veaa087

May FJ, Davis CT, Tesh RB, Barrett ADT (2011) Phylogeography of West Nile Virus: from the Cradle of Evolution in Africa to Eurasia, Australia, and the Americas. Journal of Virology 85:2964–2974. 10.1128/jvi.01963-10

McMullen AR, Albayrak H, May FJ, Davis CT, Beasley DWC, Barrett ADT (2013) Molecular evolution of lineage 2 West Nile virus. J Gen Virol 94:318–325. 10.1099/vir.0.046888-0

Mohammed MA, Galbraith SE, Radford AD, Dove W, Takasaki T, Kurane I, Solomon T (2011) Molecular phylogenetic and evolutionary analyses of Muar strain of Japanese encephalitis virus reveal it is the missing fifth genotype. Infect Genet Evol 11:855–862. 10.1016/j.meegid.2011.01.020

Moreira Salles AP, de Seixas Santos Nastri AC, Ho YL, Vilas Boas Casadio L, Emanuel Amgarten D, Justo Arevalo S, Soares Gomes-Gouvea M, Jose Carrilho F, de Mello Malta F, Rebello Pinho JR (2022) Updating the Phylodynamics of Yellow Fever Virus 2016-2019 Brazilian Outbreak With New 2018 and 2019 Sao Paulo Genomes. Front Microbiol 13:811318. 10.3389/fmicb.2022.811318

Moureau G, Cook S, Lemey P, Nougairede A, Forrester NL, Khasnatinov M, Charrel RN, Firth AE, Gould EA, de Lamballerie X (2015) New insights into flavivirus evolution, taxonomy and biogeographic history, extended by analysis of canonical and alternative coding sequences. PLoS One 10:e0117849. 10.1371/journal.pone.0117849

O’Toole A, Neher RA, Ndodo N, Borges V, Gannon B, Gomes JP, Groves N, King DJ, Maloney D, Lemey P, Lewandowski K, Loman N, Myers R, Omah IF, Suchard MA, Worobey M, Chand M, Ihekweazu C, Ulaeto D, Adetifa I, Rambaut A (2023) APOBEC3 deaminase editing in mpox virus as evidence for sustained human transmission since at least 2016. Science 382:595–600. 10.1126/science.adg8116

Pan XL, Liu H, Wang HY, Fu SH, Liu HZ, Zhang HL, Li MH, Gao XY, Wang JL, Sun XH, Lu XJ, Zhai YG, Meng WS, He Y, Wang HQ, Han N, Wei B, Wu YG, Feng Y, Yang DJ, Wang LH, Tang Q, Xia G, Kurane I, Rayner S, Liang GD (2011) Emergence of genotype I of Japanese encephalitis virus as the dominant genotype in Asia. J Virol 85:9847–9853. 10.1128/JVI.00825-11

Postler TS, Beer M, Blitvich BJ, Bukh J, de Lamballerie X, Drexler JF, Imrie A, Kapoor A, Karganova GG, Lemey P, Lohmann V, Simmonds P, Smith DB, Stapleton JT, Kuhn JH (2023) Renaming of the genus Flavivirus to Orthoflavivirus and extension of binomial species names within the family Flaviviridae. Arch Virol 168:224. 10.1007/s00705-023-05835-1

Rambaut A, Drummond AJ, Dong X, Baele G, Suchard MA (2018) Posterior summarization in Bayesian phylogenetics using tracer 1.7. Syst. Biol. 67:901–904. 10.1093/sysbio/syy032

Rambaut A, Lam TT, Max Carvalho L, Pybus OG (2016) Exploring the temporal structure of heterochronous sequences using TempEst (formerly Path-O-Gen). Virus Evol. 2:vew007. 10.1093/ve/vew007

Ravanini P, Huhtamo E, Ilaria V, Crobu MG, Nicosia AM, Servino L, Rivasi F, Allegrini S, Miglio U, Magri A, Minisini R, Vapalahti O, Boldorini R (2012) Japanese encephalitis virus RNA detected in Culex pipiens mosquitoes in Italy. Euro Surveill 17. 10.2807/ese.17.28.20221-en

Ricklin ME, Garcia-Nicolas O, Brechbuhl D, Python S, Zumkehr B, Nougairede A, Charrel RN, Posthaus H, Oevermann A, Summerfield A (2016) Vector-free transmission and persistence of Japanese encephalitis virus in pigs. Nat Commun 7:10832. 10.1038/ncomms10832

Schuh AJ, Ward MJ, Brown AJ, Barrett AD (2013) Phylogeography of Japanese encephalitis virus: genotype is associated with climate. PLoS Negl Trop Dis 7:e2411. 10.1371/journal.pntd.0002411

Sikazwe C, Neave MJ, Michie A, Mileto P, Wang J, Cooper N, Levy A, Imrie A, Baird RW, Currie BJ, Speers D, Mackenzie JS, Smith DW, Williams DT (2022) Molecular detection and characterisation of the first Japanese encephalitis virus belonging to genotype IV acquired in Australia. PLoS Negl Trop Dis 16:e0010754. 10.1371/journal.pntd.0010754

Simon-Loriere E, Faye O, Prot M, Casademont I, Fall G, Fernandez-Garcia MD, Diagne MM, Kipela JM, Fall IS, Holmes EC, Sakuntabhai A, Sall AA (2017) Autochthonous Japanese Encephalitis with Yellow Fever Coinfection in Africa. N Engl J Med 376:1483–1485. 10.1056/NEJMc1701600

Sistrom M, Andrews H, Edwards DL (2024) Comparative genomics of Japanese encephalitis virus shows low rates of recombination and a small subset of codon positions under episodic diversifying selection. PLoS Negl Trop Dis 18:e0011459. 10.1371/journal.pntd.0011459

Srihi H, Chatti N, Ben Mhadheb M, Gharbi J, Abid N (2021) Phylodynamic and phylogeographic analysis of the complete genome of the West Nile virus lineage 2 (WNV-2) in the Mediterranean basin. BMC Ecol Evol 21:183. 10.1186/s12862-021-01902-w

Srivastava KS, Jeswani V, Pal N, Bohra B, Vishwakarma V, Bapat AA, Patnaik YP, Khanna N, Shukla R (2023) Japanese Encephalitis Virus: An Update on the Potential Antivirals and Vaccines. Vaccines (Basel) 11. 10.3390/vaccines11040742

Stica CJ, Barrero RA, Murray RZ, Devine GJ, Phillips MJ, Frentiu FD (2022) Global Evolutionary History and Dynamics of Dengue Viruses Inferred from Whole Genome Sequences. Viruses 14. 10.3390/v14040703

Sumiyoshi H, Mori C, Fuke I, Morita K, Kuhara S, Kondou J, Kikuchi Y, Nagamatu H, Igarashi A (1987) Complete nucleotide sequence of the Japanese encephalitis virus genome RNA. Virology 161:497–510. 10.1016/0042-6822(87)90144-9

Sun JW, D., Zhong H, Guan D, Zhang H, Tan Q, Zhou H, Zhang M, Ning D, Zhang B, Ke C, Song T, Lin J, Zhang Y, Koopmans M, Gao GF (2017) Returning ex-patriot Chinese to Guangdong, China, increase the risk for local transmission of Zika virus. J Infect 75:356–367. 10.1016/j.jinf.2017.07.001

Takhampunya R, Kim HC, Tippayachai B, Kengluecha A, Klein TA, Lee WJ, Grieco J, Evans BP (2011) Emergence of Japanese encephalitis virus genotype V in the Republic of Korea. Virol J 8:449. 10.1186/1743-422X-8-449

Theze J, Li T, du Plessis L, Bouquet J, Kraemer MUG, Somasekar S, Yu G, de Cesare M, Balmaseda A, Kuan G, Harris E, Wu CH, Ansari MA, Bowden R, Faria NR, Yagi S, Messenger S, Brooks T, Stone M, Bloch EM, Busch M, Munoz-Medina JE, Gonzalez-Bonilla CR, Wolinsky S, Lopez S, Arias CF, Bonsall D, Chiu CY, Pybus OG (2018) Genomic Epidemiology Reconstructs the Introduction and Spread of Zika Virus in Central America and Mexico. Cell Host Microbe 23:855–864 e857. 10.1016/j.chom.2018.04.017

Van den Eynde C, Sohier C, Matthijs S, De Regge N (2022) Japanese Encephalitis Virus Interaction with Mosquitoes: A Review of Vector Competence, Vector Capacity and Mosquito Immunity. Pathogens 11. 10.3390/pathogens11030317

Vannice KS, Hills SL, Schwartz LM, Barrett AD, Heffelfinger J, Hombach J, Letson GW, Solomon T, Marfin AA, Japanese encephalitis vaccination experts p (2021) The future of Japanese encephalitis vaccination: expert recommendations for achieving and maintaining optimal JE control. NPJ Vaccines 6:82. 10.1038/s41541-021-00338-z

Williams DT, Diviney SM, Niazi AU, Durr PA, Chua BH, Herring B, Pyke A, Doggett SL, Johansen CA, Mackenzie JS (2015) The Molecular Epidemiology and Evolution of Murray Valley Encephalitis Virus: Recent Emergence of Distinct Sub-lineages of the Dominant Genotype 1. PLoS Negl Trop Dis 9:e0004240. 10.1371/journal.pntd.0004240

Xu G, Gao T, Wang Z, Zhang J, Cui B, Shen X, Zhou A, Zhang Y, Zhao J, Liu H, Liang G (2023) Re-Emerged Genotype IV of Japanese Encephalitis Virus Is the Youngest Virus in Evolution. Viruses 15. 10.3390/v15030626

Zecchin B, Fusaro A, Milani A, Schivo A, Ravagnan S, Ormelli S, Mavian C, Michelutti A, Toniolo F, Barzon L, Monne I, Capelli G (2021) The central role of Italy in the spatial spread of USUTU virus in Europe. Virus Evol 7:veab048. 10.1093/ve/veab048

